# A review of diagnostic approaches to medullary thyroid carcinoma

**DOI:** 10.1101/116905

**Authors:** Soheila Borhani, Mehdi Hedayati

## Abstract

Medullary thyroid carcinoma (MTC), occurring sporadically or as an autosomal dominant trait, accounts for 5-10% of all thyroid gland neoplasms. While the sporadic variant originates from somatic rearranged-during-transfection (RET) mutations, the inherited variant is preceded by germ-line mutations of the RET proto-oncogene. Analysis of the transformations of this certain gene is the cornerstone of diagnostic and prognostic approaches in MTC. Additionally, a panel of histopathological evaluations, biochemical markers, and imaging procedures play a pivotal role in the management of MTC. The survival rate of MTC is relatively low compared to differentiated thyroid neoplasia and is highly influenced by the stage of tumors. Improvement in MTC surveillance significantly depends on early diagnosis as well as implementation of proper screening modalities in hereditary subtypes. The present review addresses medullary thyroid carcinoma, and in particular, the current diagnostic approaches to this challenging malignancy.

## 1. Introduction

Thyroid cancer is the most prevalent endocrine malignancy, and accounts for 3.8% of all newly diagnosed cancer cases. Medullary thyroid carcinoma (MTC) originates from the parafollicular or C-cells of the thyroid, and currently constituted for 5%–10% of thyroid neoplasia. The C-cells dispersed throughout the thyroid, while they are predominantly localized to the upper two-thirds of each lobe of the gland. Medullary thyroid carcinoma mainly occurs sporadically. However, about 25% of the cases are transmitted in an autosomal dominant pattern. Hereditary MTC most often presents as part of the multiple endocrine neoplasias (MEN) type 2 syndromes, otherwise inherited as a pure familial medullary cancer (FMTC) [1–5]. The inherited subtypes of MTC are associated with the germ-line mutations of Rearranged during Transfection (RET) proto-oncogene. The RET gene encodes a receptor tyrosine kinase (RTK), that is expressed in parafollicular cell lines of the thyroid gland and the other embryonic neural crest derivatives such as chromaffin cells of adrenal and parathyroid cell lines. The characteristic distribution of RTK could explain diverse phenotypes of the MEN2 syndromes, including MTC, pheochromocytoma and/or hyperparathyroidism [6–8].

Medullary thyroid carcinoma exhibits a variety of clinical courses, from an indolent neoplasm to an aggressive tumor with high mortality. Surveillance of MTC has been discernibly compromised by increasing the patient’s age as well as the stage of the tumor based on the TNM (tumor, node, and metastasis) classification system of the American Joint Committee on Cancer (AJCC). Patients suffering from distant metastatic conditions have a considerably lower survival rate, while individuals with tumors confined to the thyroid could be completely cured. Metastasis is explored in approximately 10%–15% of patients at initial diagnosis and the average survival rate of MTC is considered around 65-89% after 5 years [9–12]. Putting together, early diagnosis as well as the implementation of a precise screening panel would considerably promote the survival rate and prognosis of medullary thyroid carcinoma.

The present review identified literature addressing the current diagnostic modalities, and the latest screening approaches to medullary thyroid carcinoma as well as an overview of the clinical manifestations and epidemiological features of this malignancy. The resources included database on MTC in Pubmed and Google Scholar. There was no limitation regarding the publication date and publication status, or language. The search strategy consisted of keyword terms related to the disease, including medullary thyroid carcinoma, multiple endocrine neoplasia syndromes, and RET proto-oncogene.

## 2. Clinical manifestations of MTC

The first step in the diagnostic process of medullary thyroid carcinoma is a thorough clinical evaluation. This particular neoplasia is most likely prominent as a single thyroid nodule during a physical examination. Generally, MTC is categorized into sporadic and hereditary components. Subsequently, the hereditary subtype is characterized as three distinct clinical settings, namely MEN2A, MEN2B, and FMTC.

The sporadic MTC constitutes about 75% of all MTC cases, with a mean age of diagnosis in the fifth to sixth decades. Chronic hypercalcemia is thought of as the modulating factor, however, there is a lack of sufficient evidence-based data [13]. Sporadic MTC typically manifests a painless solitary nodule in the thyroid gland. Cervical lymph node involvement would be detected in about 50% of patients. Besides, tumor invasion to the adjacent structures occurs in 15% of cases and distant metastasis is found in nearly 10% of individuals at the time of diagnosis. Clinical presentation of MTC sometimes correlates with hormonal secretion; these tumors might produce calcitonin, prostaglandins, serotonin and vasoactive intestinal peptide (VIP), which exert systemic symptoms such as diarrhea and flushing in some patients. Cushing's syndrome is preceding by paraneoplastic secretion of adrenocorticotropic hormone (ACTH) in a few MTC cases [14, 15].

In contrast to sporadic medullary thyroid cancer, hereditary MTC tends to be multifocal and presents at a younger age. MEN 2A or Sipple’s syndrome is the most frequent component of hereditary MTC and exhibits as a triad of MTC, pheochromocytoma, and hyperparathyroidism. The MTC is a predominant feature and occurs in approximately 95% of cases. Pheochromocytoma most often develops bilaterally, being present in nearly 30%-50% of patients, and hyperparathyroidism is usually due to parathyroid hyperplasia and comprises 10%-20% of this syndrome. Also, MEN2A syndrome may be associated with lichen amyloidosis, a cutaneous pruritic eruption, and Hirschsprung’s disease, a congenital consequence of depleting the myenteric nervous plexuses in the hindgut.

MEN 2B syndrome has a more aggressive clinical course with high mortality rate and occurs in early life compared to the other subtypes of hereditary MTC. This syndrome is characterized by MTC in 90%, pheochromocytoma in 45% and ganglioneuromatosis in almost all cases. Patients have multiple mucosal neuromas of the anterolateral surface of the tongue, lips, and conjunctivae. The ganglioneuromatosis of the gastrointestinal tract may contribute to enteric disturbances, such as intestinal obstruction or constipation. Besides, some cases of MEN 2B syndrome demonstrate musculoskeletal deformities. Occasionally, individuals with typical features of MEN 2A or MEN 2B syndrome would have any identifiable RET mutation. On the occurrence of such condition, at least two clinical characteristics of the disorders are required to reach adequate diagnostic accuracy [16–19].

Familial medullary thyroid carcinoma accounts for approximately 10% of hereditary MTC and has a relatively more favorable outcome. It is commonly presented as MTC in two or more generations of a family in the absence of certain manifestations of MEN syndromes, pheochromocytoma, and hyperparathyroidism. However, describing FMTC as an isolated syndrome is a controversial issue, and it may be defined as a variant of MEN2A syndrome with delayed clinical presentation [20, 21].

## 3. Diagnostic approaches in MTC

Detection of a thyroid nodule, particularly if concomitant with either positive family history for MTC or other endocrinopathies, would mount the probability of MTC. However, diagnosis of MTC on clinical grounds alone could be misleading. Besides, a variety of conditions may resemble medullary thyroid cancer including multinodular goiter, different forms of thyroid neoplasia or even direct invasion of neoplasms from adjacent organs to the thyroid gland [22]. The following modalities are increasingly applied in diagnosis and management of MTC: histopathologic, biochemical, imaging, and molecular analysis.

### 3.1. Histopathologic analysis

The sporadic medullary thyroid carcinoma is typically unilateral, while hereditary MTC develops in a bilateral pattern. Nevertheless, some cases of apparently sporadic MTC manifest distinctive features of hereditary variants. On gross pathologic examination, MTC appears as firm, white or gray and gritty tumor with sizes varying from small nodules to masses. The tumors are most often non-encapsulated but well demarcated. Diagnosis of MTC is mainly based on Fine-needle Aspiration Biopsy (FNAB), or Fine-needle Aspiration Cytology (FNAC). Histopathological analysis of tissues most likely reveals the salt-and-pepper–like chromatin, prominent nucleoli, and granular eosinophilic cytoplasm. The neoplastic cells are commonly arranged in sheets, trabecular, or glandular patterns. Alongside, necrosis, hemorrhage, lymphatic invasion, and amyloid clumps may be present. Stromal amyloid deposition in the absence of follicular cells is assumed as histological characteristic of MTC. The amyloid clumps can be recognized by Congo red or Papanicolaou's staining, and appear in 50%-70% of specimens. Also, Psammoma bodies and S100+ cells are detectable in some cases of MTC [23, 24].

Application of invasive procedures for diagnosis of MTC has been attenuated by using FNA. However, an inadequate sample size of lesions may lead to false negative test results. FNAC was able to explore approximately one-half of MTC in a recent meta-analysis [25]. Implementation of ancillary techniques such as ultrasound-guided fine needle aspiration and on-site evaluation could significantly improve the diagnostic accuracy of thyroid FNA specimens [26]. On the other hand, the similarity of cytologic findings in MTC and different types of thyroid cancer could make diagnosis more challenging. In such cases, ImmunoHistoChemistry (IHC) of samples for calcitonin (Ct), calcitonin gene-related peptide (CGRP), carcinoembryonic antigen (CEA), as well as detection of calcitonin in the washout fluid (FNA-Ct) would enhance the specificity of the results [27–29]. Additionally, IHC for matrix metalloproteinases (MMPs) subtypes such as MMP-9 and tissue inhibitor of matrix metalloproteinase (TIMP-2) are implicated in the evaluation of MTC. The MMPs are involved in tumor growth, angiogenesis and distant metastasis [30].

### 3.2. Biochemical analysis

MTC produces a variety of peptides and hormones including calcitonin, carcinoembryonic antigen, amyloid, somatostatin, adrenocorticotropin, vasoactive intestinal peptide, calcitonin gene-related peptide, chromogranin A and serotonin. The most prominent hormone is calcitonin, a polypeptide that regulates serum calcium and is extensively used for diagnosis of C-cells hyperplasia and MTC. Serum Ct values of above 10 pg/ml are considered to be abnormal. Regional lymph node extensions are generally associated with basal Ct levels beyond 10-40 pg/ml and metastasis correlated with Ct values of 150-400 pg/ml [31, 32]. The borderline serum calcitonin levels are re-evaluated following administration of pentagastrin or omeprazole. However, implementation of calcitonin stimulation testing in the management of MTC remains debated [33, 34]. Therefore, measurement of some certain proteins is considered to be beneficial to the diagnosis of MTC. Plasma concentrations of osteocalcin and retinol binding protein-4 (RBP-4) tend to increase in patients affected with MTC compared to the general population. Osteocalcin acts as a calcium regulatory hormone, and retinol-binding proteins (RBPs) are contributed in modulating intracellular retinoid metabolism and gene expression [35–38].

Carcinoembryonic antigen (CEA) can be applied as a relatively sensitive biochemical marker for MTC. Serum CEA levels of more than 100 ng/mL indicate a metastatic condition. Specifically, gradual rising in the CEA values accompanied by elevated Ct levels suggest a poor prognosis. Nevertheless, isolated elevation of serum CEA may be detected outside the setting of MTC, in other neoplasia or even some non-malignant conditions [39]. Hereditary MTC is associated with pheochromocytoma and primary hyperparathyroidism. In this regard, evaluation of these endocrinopathies should be considered as an indispensable part of the diagnostic approach. Elevated plasma or 24-hour urine concentrations of metanephrines and normetanephrine in pheochromocytoma is compatible with the finding of episodic hypertension in these patients. Furthermore, hyperparathyroidism becomes manifest as osteitis fibrosa cystica and morbidities consistent with hypercalcemia such as renal stones. Serum calcium (albumin-corrected or ionized), parathyroid hormone, alkaline phosphatase (ALP) values as well as urinary cyclic adenosine monophosphate (cAMP) levels are typically raised in the case of hyperparathyroidism. Rarely, MTC might present as a hypereosinophilic syndrome which entails complete blood count analysis [40–42].

### 3.3. Imaging procedures

MTC commonly demonstrates as a solitary thyroid nodule, and the neck ultrasonography would provide further details for proper diagnosis. The main ultrasonographic characteristics of MTC include hypoechogenicity of the nodule, intranodular calcifications, increased Doppler flow, irregularity of borders, and absence of halo. Additionally, ultrasound-based elastography is a novel diagnostic approach, whereby tissue stiffness is displayed in a spectrum of colors from red (soft tissue) to blue (hard tissue). Notwithstanding, ultrasonography does not fully detect lymph nodes located in the deep neck compartments. Also, optimal interpretation of the results highly depends on radiologists’ expertise [43–44]. In MTC cases with a suspicion of invasion or metastatic disease, ultrasonography has been superseded by more sensitive imaging procedures, such as three phase-contrast enhanced computed tomography (CT) and gadolinium enhanced magnetic resonance imaging (MRI). MTC tends to locally invade the lymph nodes, and hematogenous metastasis mainly spread into the liver. Taking together, patients with probable distant metastasis must undergo CT of neck and mediastinum in addition to MRI of liver. The nuclear medicine imaging procedures are implicated in the diagnosis of pheochromocytoma during management of MTC. 123I-metaiodobenzylguanidine (123I-MIBG) scintigraphy, 18F-fluoro dopamine (FDA), 68Ga-labelled peptides, and 18F-fluorodeoxyglucose (FDG) positron emission tomography are the predominantly used modalities. However, the latter has some limitations, especially in patients with calcitonin levels of less than 500 pg/ml [45–49].

### 3.4. Molecular analysis

Hereditary MTC is an autosomal dominant trait, characterized by missense mutations in the RET proto-oncogene. Approximately 95% of individuals affected with MEN 2 syndromes and 88% of FMTC cases have an identifiable germline RET mutation, and RET oncogene somatic mutations have been detected in 40%–50% of sporadic MTC. Molecular genetic studies have a crucial role in diagnostic approaches to the MTC, particularly for distinguishing between sporadic and hereditary variants [21, 50]. The RET proto-oncogene was identified by Takahashi et al [51]. This gene encodes a membrane spanning receptor tyrosine kinase (RTK). The extracellular domain of RET protein consists of four cadherin-like domains or CLDs, and a region rich in cysteine residues which are responsible for cell proliferation and differentiation. The intracellular portion of RTK contains two tyrosine-kinase domains that are involved in activation of intracellular signaling pathways. Germline RET mutations in hereditary MTC lead to a gain of function, whereby a single point mutation encompasses further malignant transformation [8, 52].

RET proto-oncogene is located on chromosome 10q11.2 and consists of 21 exons. The most frequent point mutations are detected in codons 609, 611, 618, and 620 relevant to exon 10 as well as in codons 630 and 634 within exon 11, and also the codon 918 in exon 16. The exons 10 and 11 are associated with the extracellular domain of RET protein, while exon 16 is pertaining to an intracellular portion of the receptor [53–58]. Although the role of RET proto-oncogene mutations in the development of MTC has been conclusively established, the impact of RET polymorphisms as a predisposing factor for medullary thyroid carcinoma is not yet well understood. Considering this issue, several studies investigated the frequency of RET variants such as G691S (exon 11, rs1799939), S904S (exon 15, rs1800863), S836S (exon 14, rs1800862), L769L (exon 13, rs1800861), and intron 14 (IVS14–24; rs2472737) polymorphisms in the individuals with MTC from diverse geographic regions [59–62]. Notwithstanding, there are some reports in the literature that refute the association between this particular SNP and development of MTC [63–66]. These findings necessitate further studies on the genetic profile of MTC in order to explain the discrepancies among patient populations from diverse ethnic origins. Furthermore, the role of phosphoinositide 3-kinase-protein kinase B/AKT (PI3K-PKB/AKT) has been investigated in the pathogenesis of thyroid cancer. This particular signaling pathway plays a key impact in multiple cellular processes, and its continuous activation by several aberrant RTKs would result in high cellular proliferation and a number of malignancies including MTC [67].

## 4. Genotype-phenotype interactions in MTC

The mutations in RET proto-oncogene harbor specific clinical manifestations. In MEN2A syndrome, the most prevalent point mutations occur in codons 609, 611, 618, 620, 630, and particularly in codon 634. Mutations of codon 634 are associated with other comorbidities including pheochromocytoma, hyperparathyroidism, and lichen amyloidosis. Hirschsprung's disease (HD) is linked to certain mutations in codon 620 leading to inactivation of the encoded protein. Some activating mutations have been evidenced as precursors of HD [68, 69]. The specific clinical features of MEN2A syndrome can be influenced by patient's genetic characteristics. Ghazi et al. reported a case of MTC with the latent presentation of pheochromocytoma as well as hyperparathyroidism. Besides, molecular analysis of the patient and her family members revealed the C634R mutation within codon 11 and a few SNPs including G691S, S836S, and S904S associated with codons 11, 14, and 15, respectively. This specific genetic profile might explain the indolent clinical course of MEN2A syndrome in this case [70]. In the majority of patients with familial medullary thyroid carcinoma, RET mutations are verified within exon 10 and exon 11, similar to those recognized in MEN2A syndrome. Moreover, point mutations in codon 768 (exon 13), codon 804 (exon 14) and codon 891 (exon 15) are also detected. In approximately 95% of patients manifesting as MEN2B syndrome, mutation M918T within exon 16 is present, and in some cases, a double mutation V804M/Y806C in exon 14 have been identified. The sporadic MTC is occasionally associated with somatic RET mutations, mostly in codons 918 (exon 16), 618, 620, 634, and 883 [71–73].

## 5. Screening approaches in MTC

The specific mutations in RET proto-oncogene not only associated with the clinical course but also would be helpful in predicting the progression of medullary thyroid carcinoma. The American Thyroid Association (ATA) described four levels of risks in evaluating the aggressiveness of these tumors. The risk level D involves point mutations in codons 883, 918 and some certain dual mutations. Patients with this set of RET mutations are at highest risk of developing MTC. Individuals with mutations in codon 634 are categorized as the level C, while the risk level B is associated with mutations in codons 609, 611,618, 620 and 630. Lastly, patients with mutations in codons 768, 790, 791, 804 and 891 are classified as level A and carry the least risk of developing and progression of MTC (Table.1).

**Table 1.**
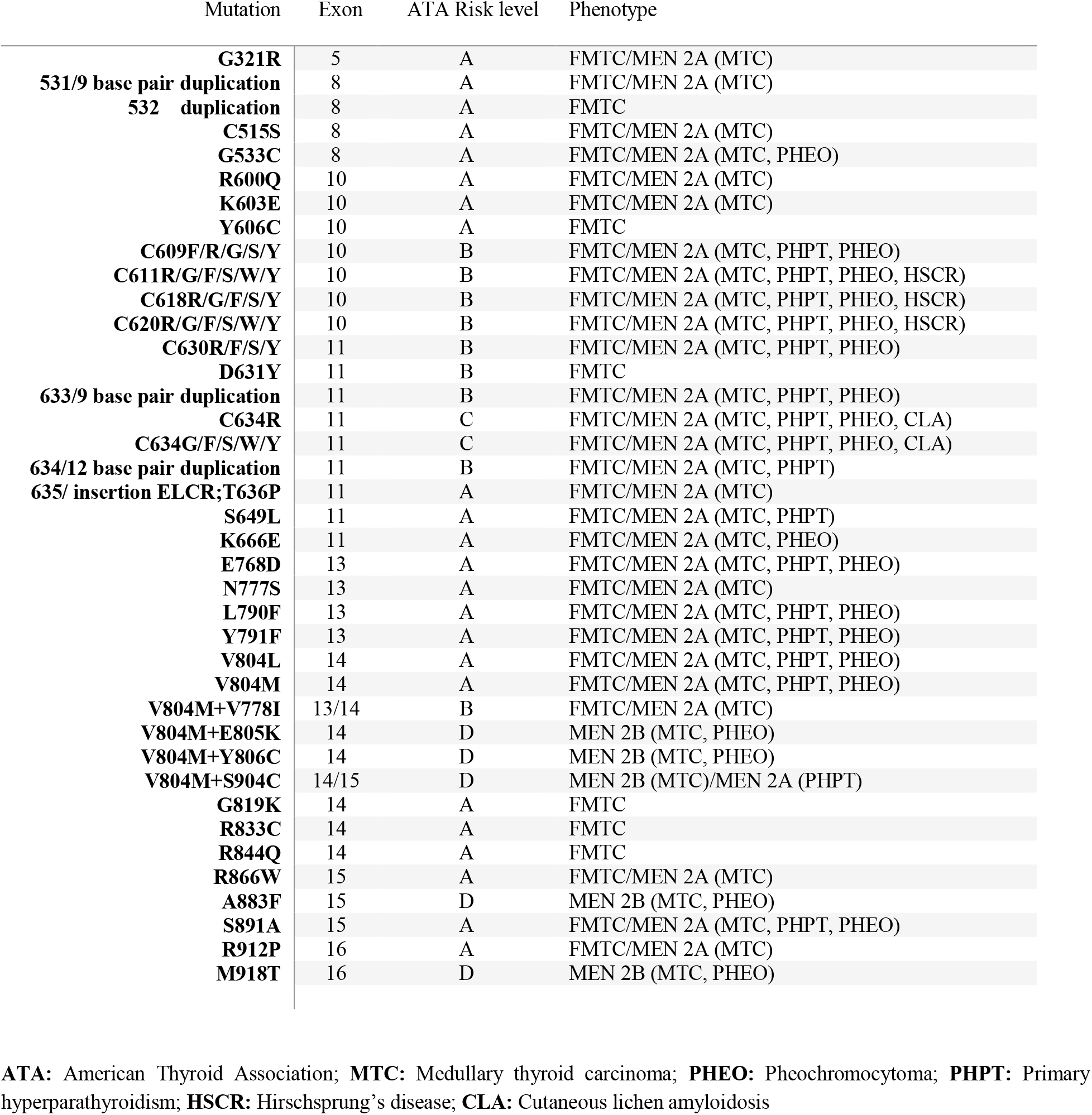
Genotype-Phenotype interactions in MTC and risk stratification according to the American Thyroid Association [19].

| Mutation | Exon | ATA Risk level | Phenotype |
| --- | --- | --- | --- |
| **G321R** | 5 | A | FMTC/MEN 2A (MTC) |
| **531/9 base pair duplication** | 8 | A | FMTC/MEN 2A (MTC) |
| **532 duplication** | 8 | A | FMTC |
| **C515S** | 8 | A | FMTC/MEN 2A (MTC) |
| **G533C** | 8 | A | FMTC/MEN 2A (MTC, PHEO) |
| **R600Q** | 10 | A | FMTC/MEN 2A (MTC) |
| **K603E** | 10 | A | FMTC/MEN 2A (MTC) |
| **Y606C** | 10 | A | FMTC |
| **C609F/R/G/S/Y** | 10 | B | FMTC/MEN 2A (MTC, PHPT, PHEO) |
| **C611R/G/F/S/W/Y** | 10 | B | FMTC/MEN 2A (MTC, PHPT, PHEO, HSCR) |
| **C618R/G/F/S/Y** | 10 | B | FMTC/MEN 2A (MTC, PHPT, PHEO, HSCR) |
| **C620R/G/F/S/W/Y** | 10 | B | FMTC/MEN 2A (MTC, PHPT, PHEO, HSCR) |
| **C630R/F/S/Y** | 11 | B | FMTC/MEN 2A (MTC, PHPT, PHEO) |
| **D631Y** | 11 | B | FMTC |
| **633/9 base pair duplication** | 11 | B | FMTC/MEN 2A (MTC, PHPT, PHEO) |
| **C634R** | 11 | C | FMTC/MEN 2A (MTC, PHPT, PHEO, CLA) |
| **C634G/F/S/W/Y** | 11 | C | FMTC/MEN 2A (MTC, PHPT, PHEO, CLA) |
| **634/12 base pair duplication** | 11 | B | FMTC/MEN 2A (MTC, PHPT) |
| **635/ insertion ELCR;T636P** | 11 | A | FMTC/MEN 2A (MTC) |
| **S649L** | 11 | A | FMTC/MEN 2A (MTC, PHPT) |
| **K666E** | 11 | A | FMTC/MEN 2A (MTC, PHEO) |
| **E768D** | 13 | A | FMTC/MEN 2A (MTC, PHPT, PHEO) |
| **N777S** | 13 | A | FMTC/MEN 2A (MTC) |
| **L790F** | 13 | A | FMTC/MEN 2A (MTC, PHPT, PHEO) |
| **Y791F** | 13 | A | FMTC/MEN 2A (MTC, PHPT, PHEO) |
| **V804L** | 14 | A | FMTC/MEN 2A (MTC, PHPT, PHEO) |
| **V804M** | 14 | A | FMTC/MEN 2A (MTC, PHPT, PHEO) |
| **V804M+V778I** | 13/14 | B | FMTC/MEN 2A (MTC) |
| **V804M+E805K** | 14 | D | MEN 2B (MTC, PHEO) |
| **V804M+Y806C** | 14 | D | MEN 2B (MTC, PHEO) |
| **V804M+S904C** | 14/15 | D | MEN 2B (MTC)/MEN 2A (PHPT) |
| **G819K** | 14 | A | FMTC |
| **R833C** | 14 | A | FMTC |
| **R844Q** | 14 | A | FMTC |
| **R866W** | 15 | A | FMTC/MEN 2A (MTC) |
| **A883F** | 15 | D | MEN 2B (MTC, PHEO) |
| **S891A** | 15 | A | FMTC/MEN 2A (MTC, PHPT, PHEO) |
| **R912P** | 16 | A | FMTC/MEN 2A (MTC) |
| **M918T** | 16 | D | MEN 2B (MTC, PHEO) |

**ATA:** American Thyroid Association; **MTC:** Medullary thyroid carcinoma; **PHEO:** Pheochromocytoma; **PHPT:** Primary hyperparathyroidism; **HSCR:** Hirschsprung's disease; **CLA:** Cutaneous lichen amyloidosis

Individuals affected with C-cell hyperplasia or MTC are evaluated for the RET proto-oncogene mutations, and family members of those with established mutations must undergo a periodically screening process at 1–3-year intervals. Examination of genetically susceptible individuals should continue until the age of 50 or up to 20 years after the oldest age of initial diagnosis in the family, whichever came later. The screening approach follows the criteria regarding aggressiveness of tumors. At-risk relatives of patients with risk level D should be evaluated for RET mutations within the first year of their life, and genetic testing of individuals in other categories should be performed before the age of 3-5. Moreover, pre-implantation and antenatal genetic analysis are implemented in the prenatal assessment of suspected cases. Molecular analysis is initially applied for detection of the most common missense mutations in RET proto-oncogene. Those with no apparent nucleotide substitution have an estimated risk of 0.18% for developing hereditary MTC. Furthermore, susceptible individuals are evaluated through imaging and biochemical screening approaches. The age of required first neck ultrasonography is identical to the aforementioned RET testing schedules. Besides, the assessment of serum calcitonin is commenced by age 6 months with the risk level D, otherwise by age less than 3-5 years for the remaining risks levels [19, 74].

## 6. Conclusion

The survival rate of medullary thyroid carcinoma is relatively low compared to the other subtypes of thyroid cancer, and early diagnosis would considerably improve the surveillance. A proper diagnostic approach to MTC generally mandates molecular analysis, histopathological evaluation, biochemical assays, as well as the application of imaging procedures. Analysis of the RET proto-oncogene is regarded as the cornerstone of the management of this malignancy. Besides, genetic assays play a pivotal role in screening and prognostic approaches to the hereditary subtypes of MTC. However, implementation of optimal diagnostic modalities needs further evaluations.

